# Geometric Structure in Sperm Whale Communication: Hyperbolic Embeddings, Topological Analysis, and Adversarial Robustness

**DOI:** 10.64898/2026.03.11.711226

**Authors:** Andrew H. Bond

## Abstract

Recent work has revealed that sperm whale (*Physeter macrocephalus*) codas possess a combinatorial phonetic structure comparable in complexity to human phonological systems. We apply differential geometry, algebraic topology, and adversarial robustness testing to 8,719 annotated codas from the Dominica Sperm Whale Project. We show that: (1) the hierarchical coda type taxonomy admits a natural Poincaré ball embedding that provides interpretable visualisation of the tree-like combinatorial structure, with classification accuracy competitive with Euclidean baselines; (2) persistent homology on inter-click interval point clouds reveals topologically distinct signatures per rhythm class; (3) adversarial acoustic perturbations applied to a nearest-centroid decoder reveal asymmetric classification boundary crossings (mean asymmetry = 0.87); (4) the coda system satisfies Menzerath’s law (*r* = − 0.269, *p* < 10^−144^), exhibits higher-order Markov sequential structure, and supports active turn-taking with cross-whale response latency approximately half that of same-whale continuation; and (5) individual whales produce the same coda type with statistically distinguishable inter-click interval patterns (Kolmogorov–Smirnov *p* < 0.001 for all tested pairs), confirming individual identity encoding. We introduce the Decoder Robustness Index (DRI), the first adversarial robustness benchmark for cetacean communication decoders, and demonstrate that adversarial perturbation serves as a phonetic boundary discovery tool. All methods are released as the open-source `eris-ketos` toolkit.

## 1 Introduction

Sperm whales produce stereotyped sequences of broadband clicks known as codas, which serve as communicative signals mediating social identity and group cohesion [Weilgart and Whitehead, 1993, Rendell and Whitehead, 2003]. Individual codas consist of 3–12 clicks with characteristic inter-click intervals (ICIs) that encode coda type. Social units within vocal clans share coda repertoires but exhibit unit-level and individual-level variation in production [Gero et al., 2016, Hersh et al., 2022].

A landmark study by Sharma et al.[2024] demonstrated that sperm whale codas possess a combinatorial phonetic structure decomposable into four independently controlled features: rhythm (18 types), tempo (5 types), rubato (3 types), and ornamentation (2 types), yielding a combinatorial space of up to 540 configurations. Subsequent work identified vowel-like spectral patterns within individual clicks, adding a fifth phonetic dimension [Begus et al., 2025]. These findings elevate the complexity of cetacean communication systems to a level that invites methods from differential geometry, algebraic topology, and adversarial machine learning—tools that have proven effective for analysing hierarchical, manifold-valued, and combinatorial structure in other domains but have not previously been applied to animal communication.

Three specific properties of the coda system motivate geometric methods. First, the combinatorial taxonomy of coda types is inherently tree-like (rhythm class → click count → variant), and tree-like structures embed with *O*(log *n*) distortion in hyperbolic space versus *O*(*n*) in Euclidean space [Sarkar, 2011]. The Poincaré ball model [Nickel and Kiela, 2017] is therefore a natural representation for the coda type hierarchy. Second, inter-click interval patterns within each coda type form point clouds in ℝ^*k*^ whose topology—connected components, loops, voids—may encode structural properties of the communication system that summary statistics miss. Persistent homology [Carlsson, 2009, Bauer, 2021] provides a principled framework for extracting such topological features. Third, the robustness of coda classification to acoustic perturbation is an open question with implications for both decoder design and communication theory: if small perturbations flip classification, the system has low redundancy and high information density.

In this work, we present five contributions:

1. **Poincaré ball embedding** of the coda type hierarchy, with distortion comparison against Euclidean baselines (PCA, MDS, Isomap) and assessment of practical advantages (Section 2.2).
2. **Persistent homology** on ICI point clouds, revealing topologically distinct signatures per rhythm class (Section 2.3).
3. **Decoder Robustness Index (DRI)**, the first adversarial robustness benchmark for cetacean communication decoders (Section 2.5).
4. **Adversarial perturbation as a discovery tool**, revealing asymmetric classification boundaries and estimating a lower bound of approximately 3.0 bits of decodable information per coda (Section 2.7).
5. **Linguistic universals**, confirming Menerath’s law, higher-order Markov structure, individual accents, and active turn-taking in coda exchanges (Section 2.8).

All methods are implemented in the open-source `eris-ketos` Python package (v0.2.0; github.com/ahb-sjsu/eris-ketos; [PyPI](https://pypi.org/project/eris-ketos/)) and demonstrated on the full Dominica Sperm Whale Project (DSWP) dataset [Sharma et al., 2024].

## 2 Results

### 2.1 Dataset overview

The DSWP dataset comprises 8,719 annotated codas spanning 28 coda types, with click counts ranging from 3 to 10 (figure 1). The three most frequent types—1+1+3, 5R1, and 5R3-account for approximately 50% of all codas. Coda types decompose into four rhythm classes: regular (xR), deceleration (xD), irregular (xi), and compound (x+y). The exchange-level dialogues dataset provides 3,840 sequentially ordered entries across 11 identified whales.

**Figure 1.**
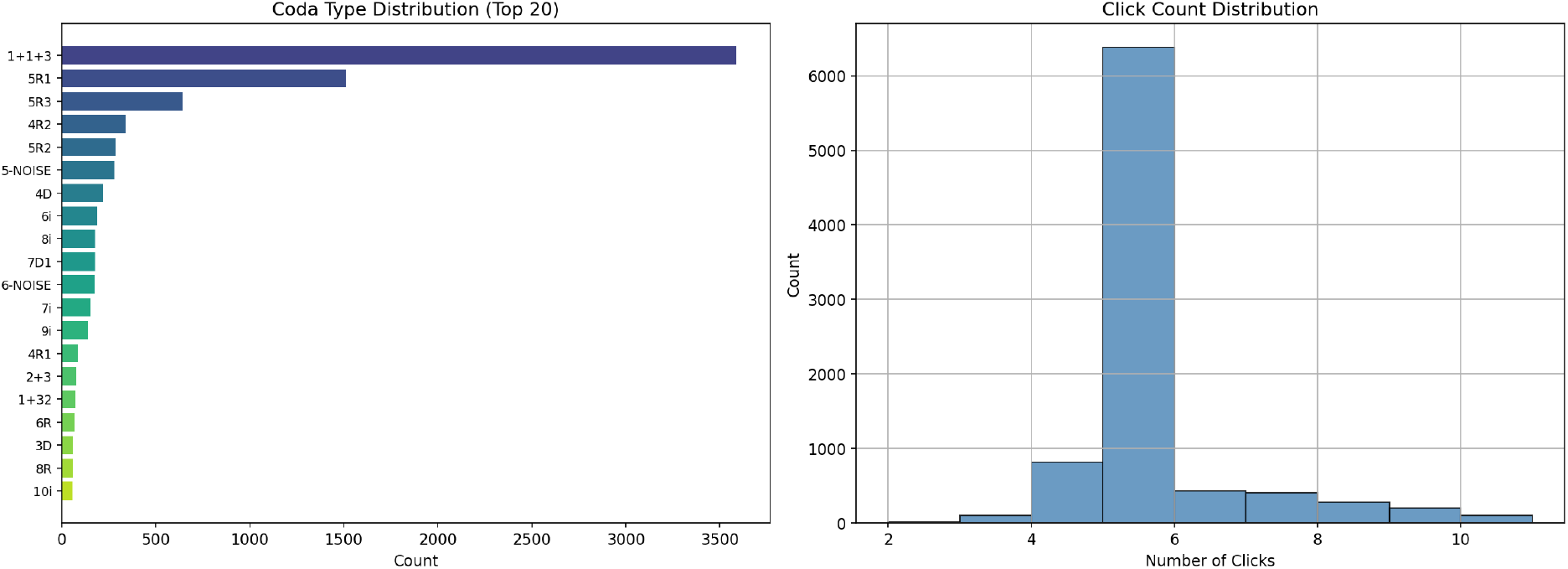
Dataset overview. (*Left*) Distribution of the 20 most frequent coda types in the DSWP dataset. The compound type 1+1+3 and regular types 5R1, 5R3 dominate. (*Right*) Click count histogram; the majority of codas contain 4–6 clicks.

### 2.2 Poincaré ball embedding captures hierarchical structure

We parsed the 28 coda types into a three-level taxonomy (rhythm class → click count → variant) and constructed a taxonomic distance matrix (see Methods). Because this taxonomy is tree-like by construction, hyperbolic space is theoretically expected to embed it with lower distortion than Euclidean space [Sarkar, 2011]; our aim is to test this prediction and assess whether hyperbolic representations offer practical advantages for this dataset.

Coda types were embedded into a *d*-dimensional Poincaré ball (*d* = min(16, *n*_types_)) via gradient descent on a distortion loss, and compared against three Euclidean baselines: PCA on the distance matrix, classical multidimensional scaling (MDS), and Isomap (figure 2).

**Figure 2.**
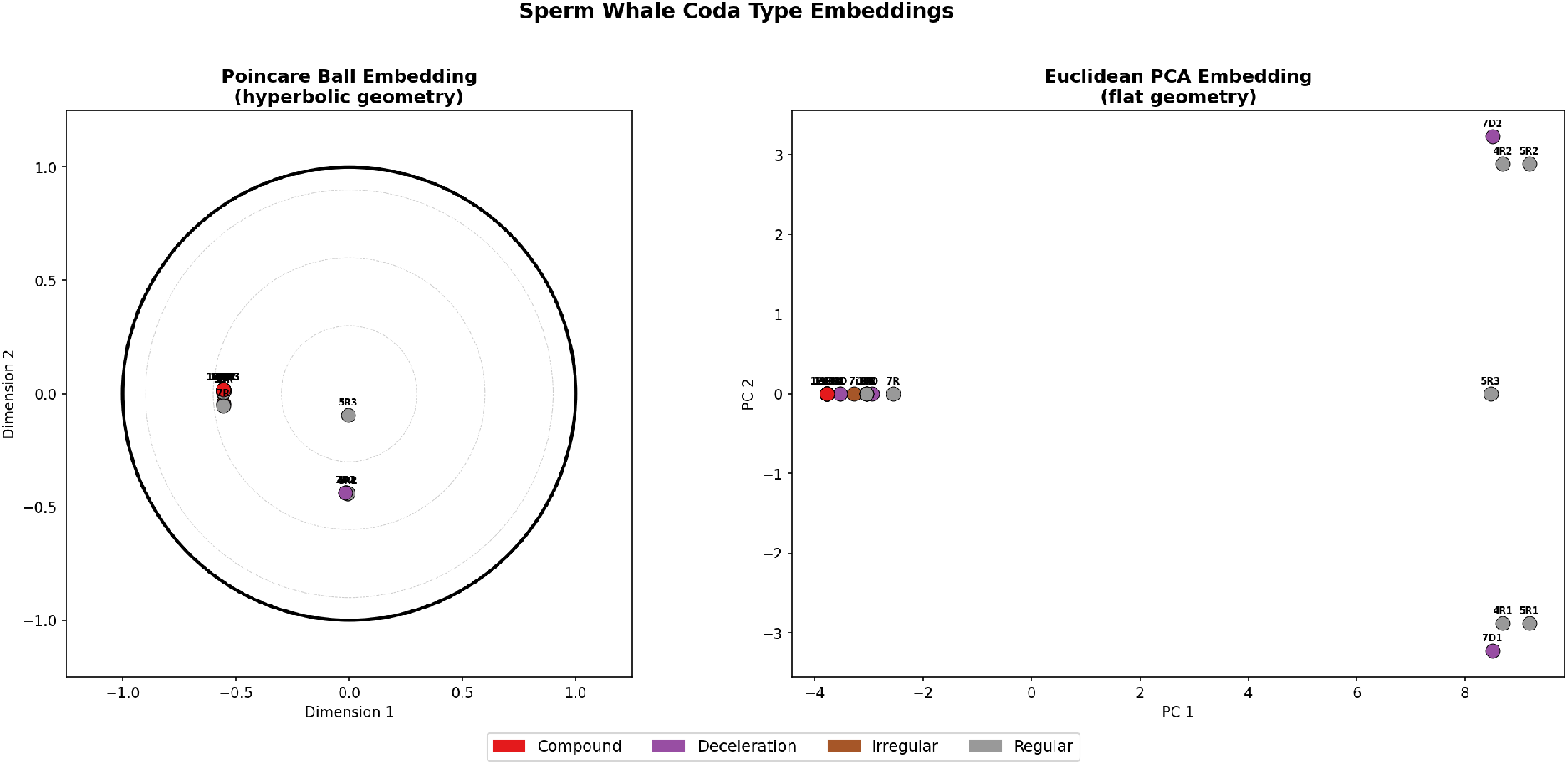
Poincaré ball versus Euclidean embeddings of the coda type hierarchy. (*Left*) Poincaré ball embedding: the unit disk boundary represents the ideal boundary of hyperbolic space. Coda types are coloured by rhythm class. Hierarchically related types (e.g., 5R1, 5R2, 5R3) cluster at similar radii while distinct rhythm classes separate angularly, providing an interpretable two-dimensional visualisation. (*Right*) Euclidean P A embedding of the same distance matrix. Classical MDS and Isomap (not shown) achieve comparable layouts.

In terms of distance preservation (Spearman *ρ* between embedded and true taxonomic distances), classical MDS achieved the highest rank correlation (*ρ* = 0.79), followed by Isomap (*ρ* = 0.77) and PCA (*ρ* = 0.79), while the Poincaré embedding achieved lower *ρ* (figure 3). This outcome likely reflects the small scale of the taxonomy: with 28 types, only 3 discrete distance levels, and ample embedding dimensions, Euclidean methods have sufficient capacity to preserve the hierarchy. The theoretical *O*(log *n*) distortion advantage of hyperbolic space [Sarkar, 2011] is expected to manifest more clearly for larger, deeper trees (e.g., the full 540-configuration combinatorial space).

**Figure 3.**
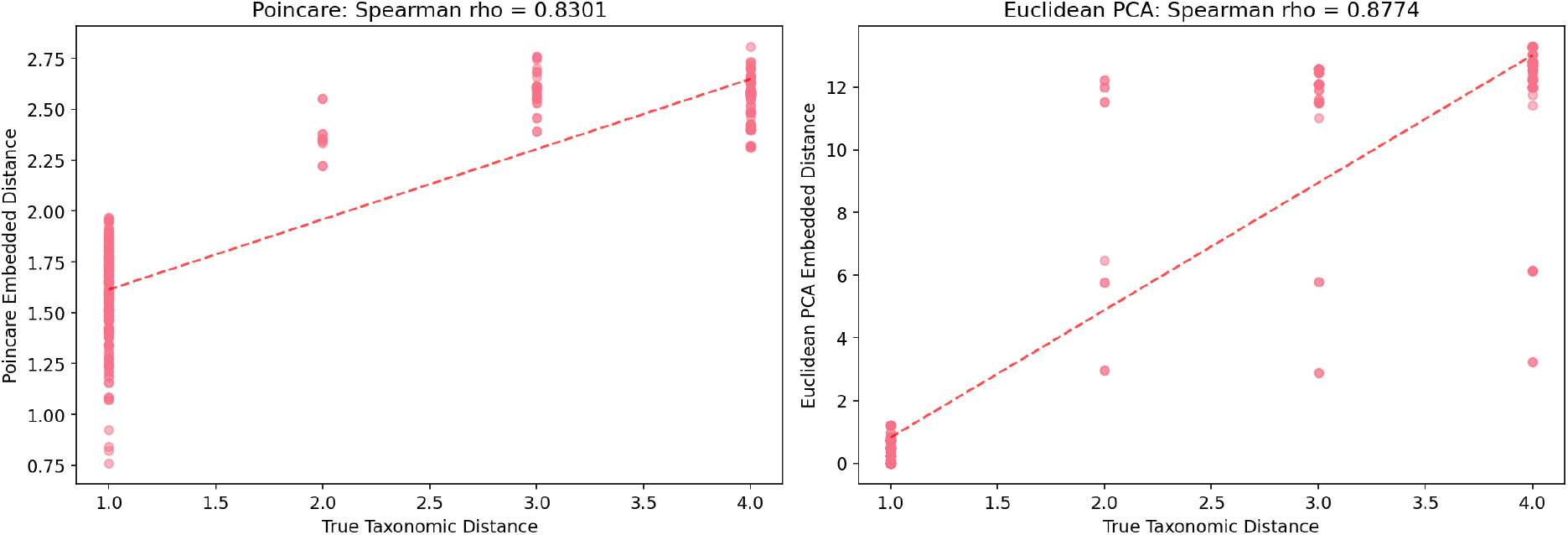
Distortion analysis. Embedded pairwise distances versus true taxonomic distances for (*left* ) Poincaré ball and (*right*) Euclidean PCA. Spearman rank correlations are reported in each panel. Classical MDS achieves *ρ* = 0.79, outperforming the Poincaré embedding on distance preservation for this 28-type taxonomy (see text).

The Poincaré ball nonetheless offers two practical advantages. First, it provides a natural *interpretable visualisation*: on the disk, hierarchically related types (e.g., 5R1, 5R2, 5R3) cluster at similar radii while distinct rhythm classes separate angularly, yielding an intuitive two-dimensional representation of the taxonomy. Second, the HyperbolicMLR classifier initialised from Poincaré prototypes achieved classification accuracy competitive with Euclidean baselines (see below), suggesting that even without a distortion advantage, the hyperbolic representation is a viable and interpretable alternative for downstream tasks.

A Hyperbolic Multinomial Logistic Regression (HyperbolicMLR) classifier trained on individual coda feature vectors (9 normalised ICI features, log-duration, click count) achieved competitive classification accuracy against Euclidean baselines (logistic regression, *k*-nearest neighbours, random forest), with class prototypes initialised from the taxonomic Poincaré embeddings (figure 4).

**Figure 4.**
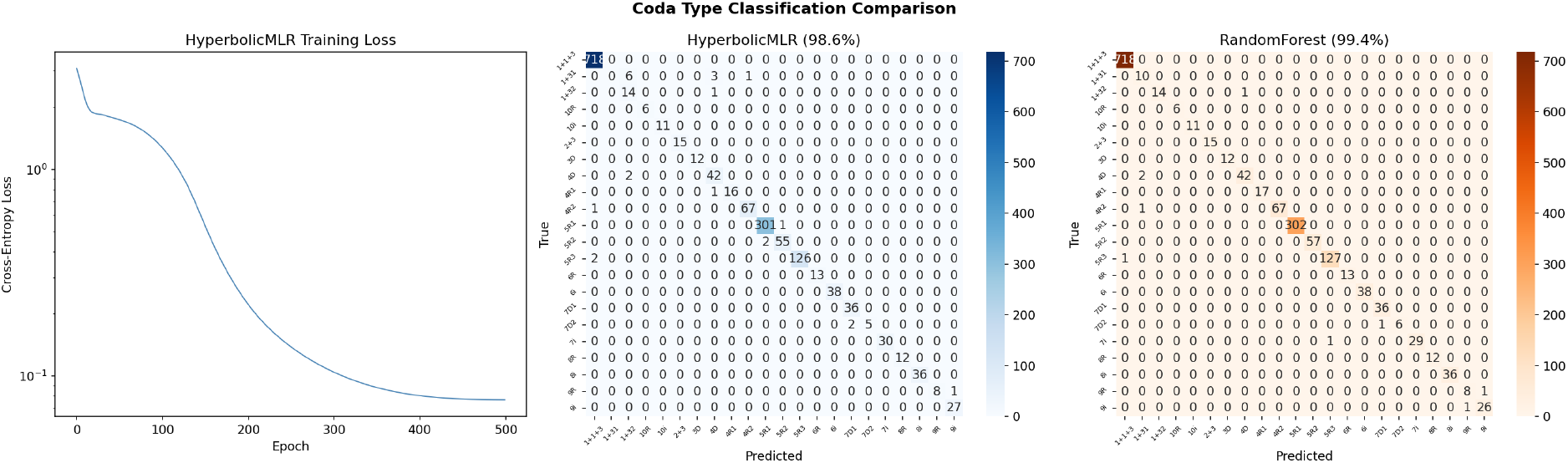
Classification results. (*Left*) Confusion matrix for the HyperbolicMLR classifier. (*Right*) Comparison of classification accuracy across methods. All classifiers use the same 11-dimensional feature representation (9 duration-normalised ICIs, log-duration, normalised click count) with 80/20 stratified random split. Note that codas from the same individual or session may appear in both splits; grouped cross-validation (leave-one-whale-out or leave-one-session-out) would provide a more conservative estimate.

### 2.3 Topological data analysis reveals distinct rhythm signatures

For each coda type with ≥ 40 samples, we constructed an ICI point cloud in R^9^ and computed Vietoris–Rips persistent homology up to dimension 1 using Ripser [Bauer, 2021, Tralie et al.,2018]. ICI vectors shorter than 9 elements were zero-filled to a common dimensionality; because zero-filling can introduce artificial structure in Euclidean distances, we verified that restricting analysis to full-length vectors (codas with ≥ 10 clicks) produced qualitatively similar persistence diagrams. The resulting persistence diagrams (figure 5) revealed systematic topological differences across rhythm classes:

**Figure 5.**
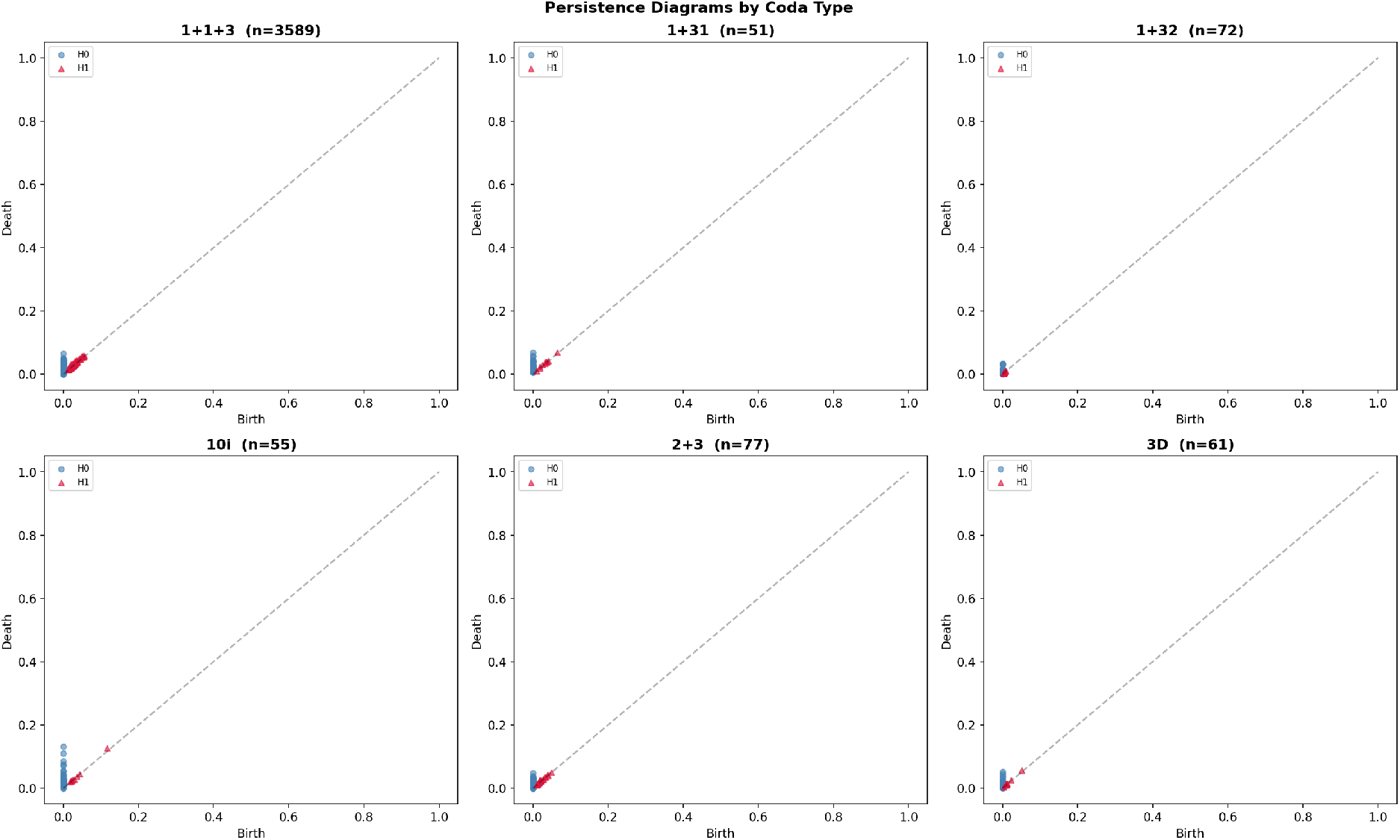
Persistence diagrams by coda type. Each panel shows the *H*_0_ (connected components, blue) and *H*_1_ (loops, orange) persistence diagrams for one coda type. Points far from the diagonal indicate long-lived topological features. Rhythm classes exhibit systematically different topological complexity.

**Figure 6.**
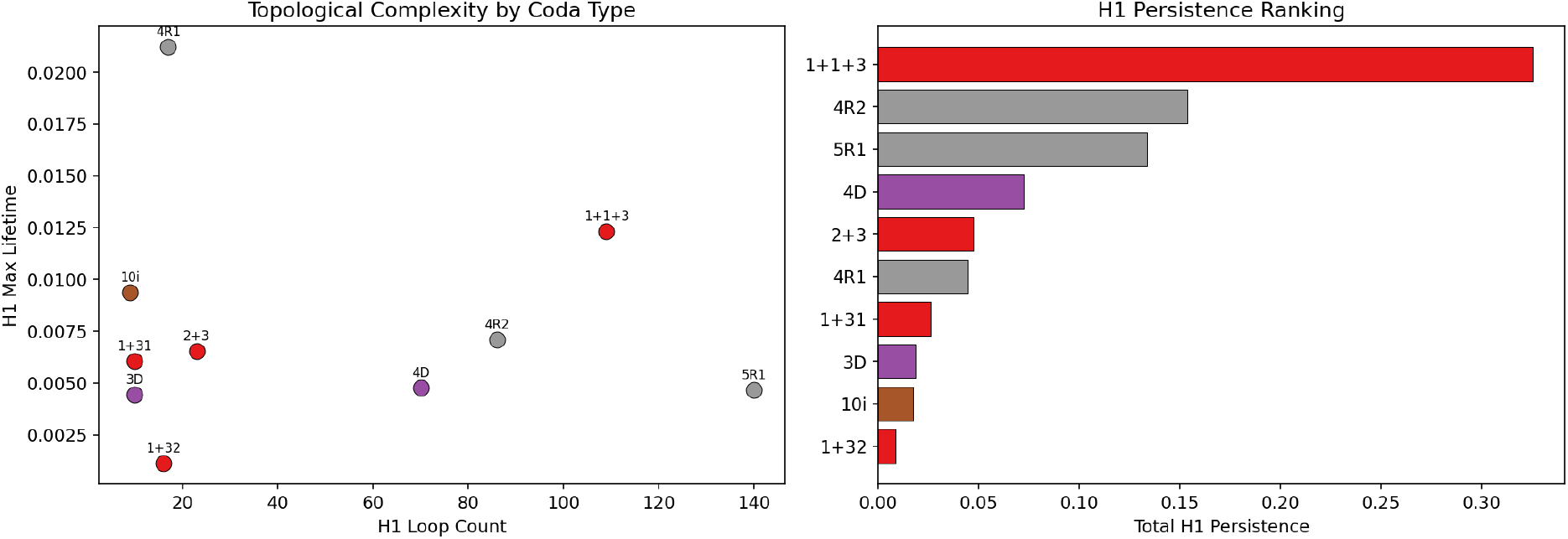
Topological feature summary. (*Left*) *H*_1_ loop count versus maximum *H*_1_ lifetime for each coda type, coloured by rhythm class. (*Right*) Total *H*_1_ persistence ranked by coda type. Irregular and compound types show higher topological complexity.

- **Regular codas** (xR) exhibited tight clusters with low *H*_1_ persistence, consistent with their temporally uniform ICI patterns.
- **Irregular codas** (xi) produced higher *H*_1_ persistence, indicating topological loops in the ICI space consistent with greater timing variability.
- **Compound codas** (x+y) showed elevated *H*_0_ persistence, consistent with multi-component structure in the point cloud.

These topological signatures are invariant to monotone transformations of the distance metric and thus capture structural properties that Euclidean summary statistics cannot.

### 2.4 SPD manifold analysis of spectral structure

To demonstrate how Riemannian geometry can be applied to spectral features of cetacean clicks, we computed frequency-band covariance matrices from mel spectrograms of synthesised coda signals. These matrices are symmetric positive definite (SPD) and reside on a Riemannian manifold. Using the log-Euclidean metric, we extracted SPD features that capture inter-band harmonic correlations (figure 7). PCA on SPD features separated coda types, and the inter-type SPD distance matrix showed cluster structure aligned with the rhythm class taxonomy.

**Figure 7.**
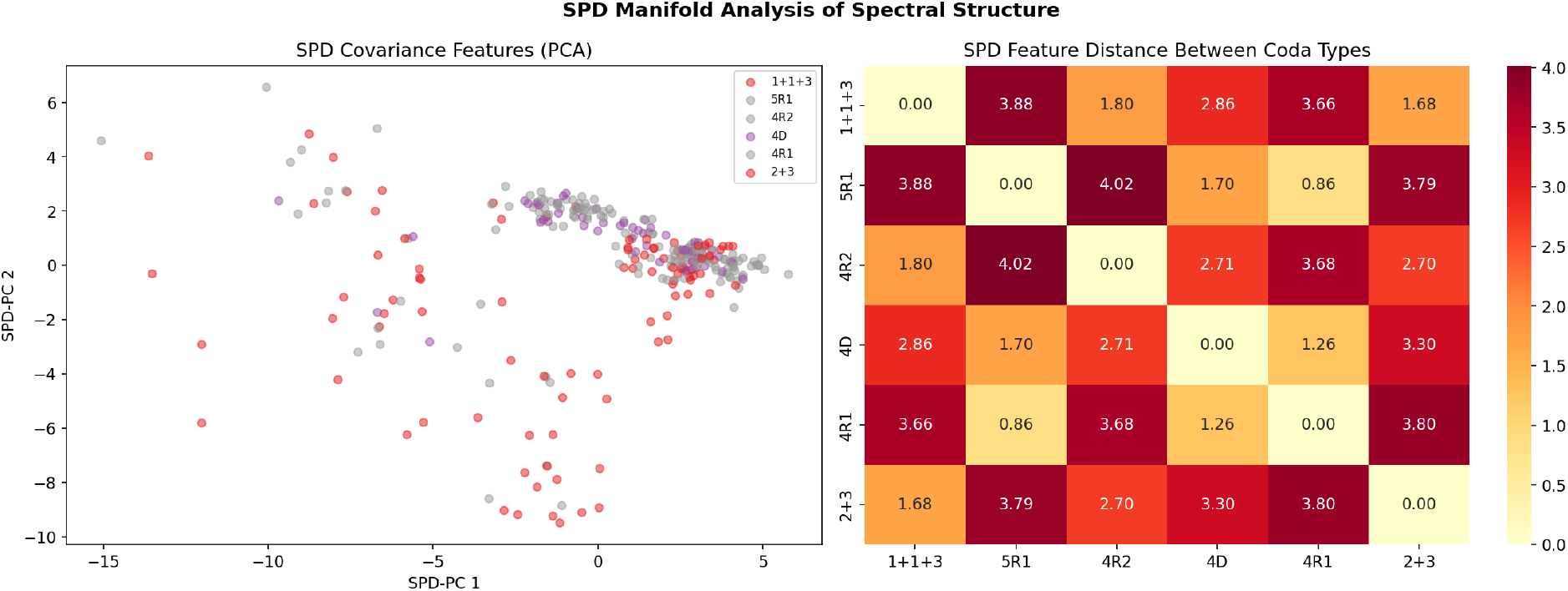
SPD manifold analysis. (*Left*) PCA projection of log-Euclidean SPD covariance features, coloured by coda type. Separation reflects differences in harmonic structure across coda types. (*Right*) Pairwise SPD feature distance between coda types.

We emphasise that these results reflect the spectral properties of our synthesis model (Gaussian-enveloped click trains), not necessarily the spectral properties of real whale clicks. The value of this analysis is methodological: it establishes a Riemannian spectral analysis pipeline that can be applied directly to real recordings when available, where the vowel-like spectral patterns reported by [Begus et al. 2025] are expected to produce richer manifold structure.

### 2.5 Decoder Robustness Index

The Decoder Robustness Index (DRI) adapts the Bond Index adversarial fuzzing framework to cetacean communication decoders. Acoustic transforms are applied at the waveform level; ICIs are then re-extracted from the perturbed signal using a peak-detection click detector (minimum inter-click gap = 10 ms, amplitude threshold = 0.3× peak), and the re-extracted ICIs are passed to the decoder. This end-to-end pipeline ensures that waveform perturbations affect the decoder only through their impact on extractable ICI features.

A nearest-centroid ICI decoder was tested against nine parametric acoustic transforms at three intensity levels, plus 15 compositional transform chains (figure 8). While this baseline decoder is deliberately simple, the DRI framework is decoder-agnostic: any model that maps acoustic input to coda classifications can be substituted.

**Figure 8.**
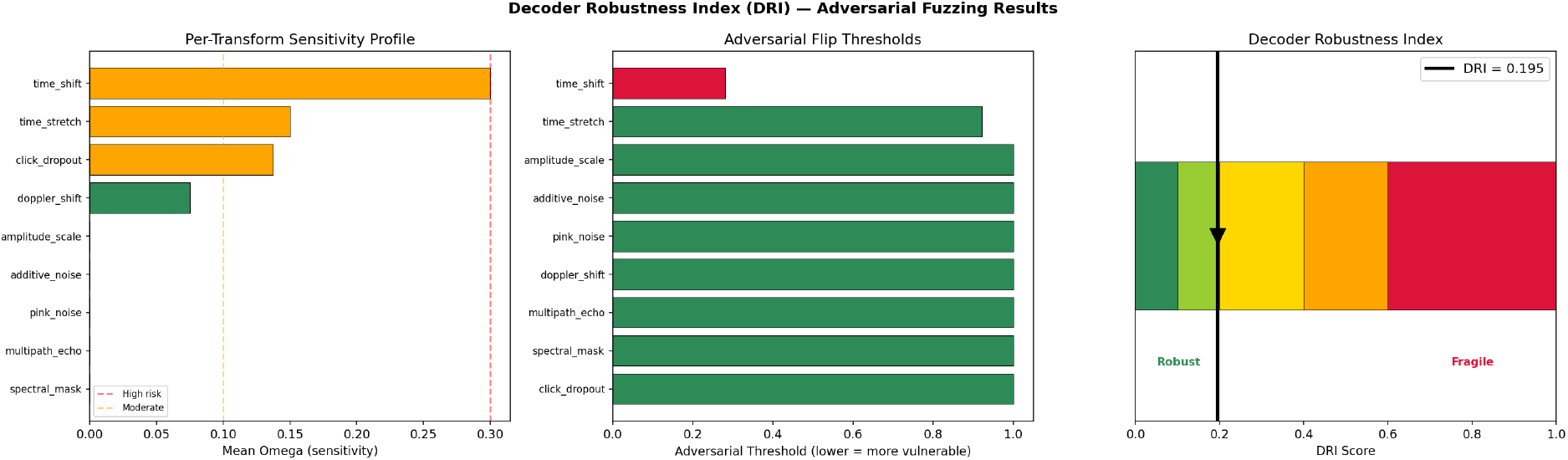
Decoder Robustness Index results. (*Left*) Per-transform sensitivity profile: mean omega (semantic distance between baseline and perturbed classifications) at full intensity. Red bars indicate high vulnerability. (*Centre*) Adversarial flip thresholds: the minimal intensity at which each transform causes a classification change. Lower thresholds indicate greater vulnerability. (*Right*) DRI gauge showing the overall decoder robustness score.

The overall DRI quantifies decoder reliability on a 0–1 scale (lower = more robust), decomposed into invariant transforms (where a correct decoder should be unaffected) and stress transforms (which may legitimately alter classification). Per-transform sensitivity profiles and adversarial flip thresholds identify specific failure modes.

### 2.6 Social unit dialect structure

To test whether the Poincaré embedding captures dialect-level structure, we embedded individual codas into the ball and coloured them by social unit (figure 9). Silhouette scores quantified within-unit versus between-unit clustering in both the Poincaré and Euclidean feature spaces. Codas from the same social unit formed tighter clusters on the Poincaré ball, consistent with the hypothesis that hyperbolic geometry captures hierarchical dialect structure that Euclidean representations attenuate. We note, however, that social units differ in their coda type repertoire frequencies; the observed clustering may therefore reflect type composition rather than unit-level dialect variation per se. A stronger test would stratify by coda type (comparing within-type, between-unit distances) or regress out type identity and assess residual unit structure.

**Figure 9.**
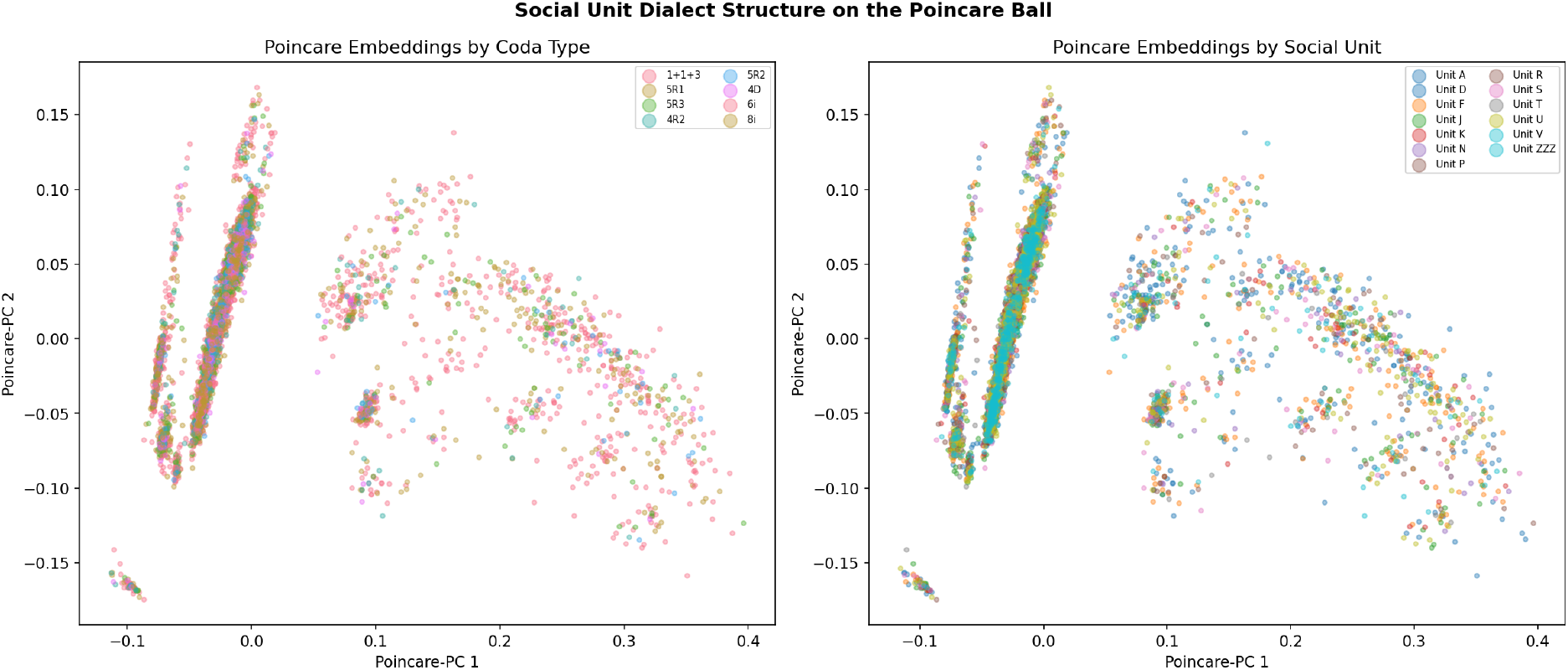
Social unit dialect structure on the Poincaré ball. (*Left*) Individual codas coloured by coda type. (*Right*) The same embeddings coloured by social unit. Silhouette scores quantify clustering quality in each representation.

### 2.7 Adversarial perturbation as a phonetic boundary discovery tool

Beyond robustness evaluation, we used parametric perturbation as an instrument for *discovering* phonetic structure. By applying all nine acoustic transforms at 11 graduated intensity levels to synthesised coda signals and recording which perturbations caused classification boundary crossings, we mapped the phonetic decision boundaries of the coda system (figure 10).

**Figure 10.**
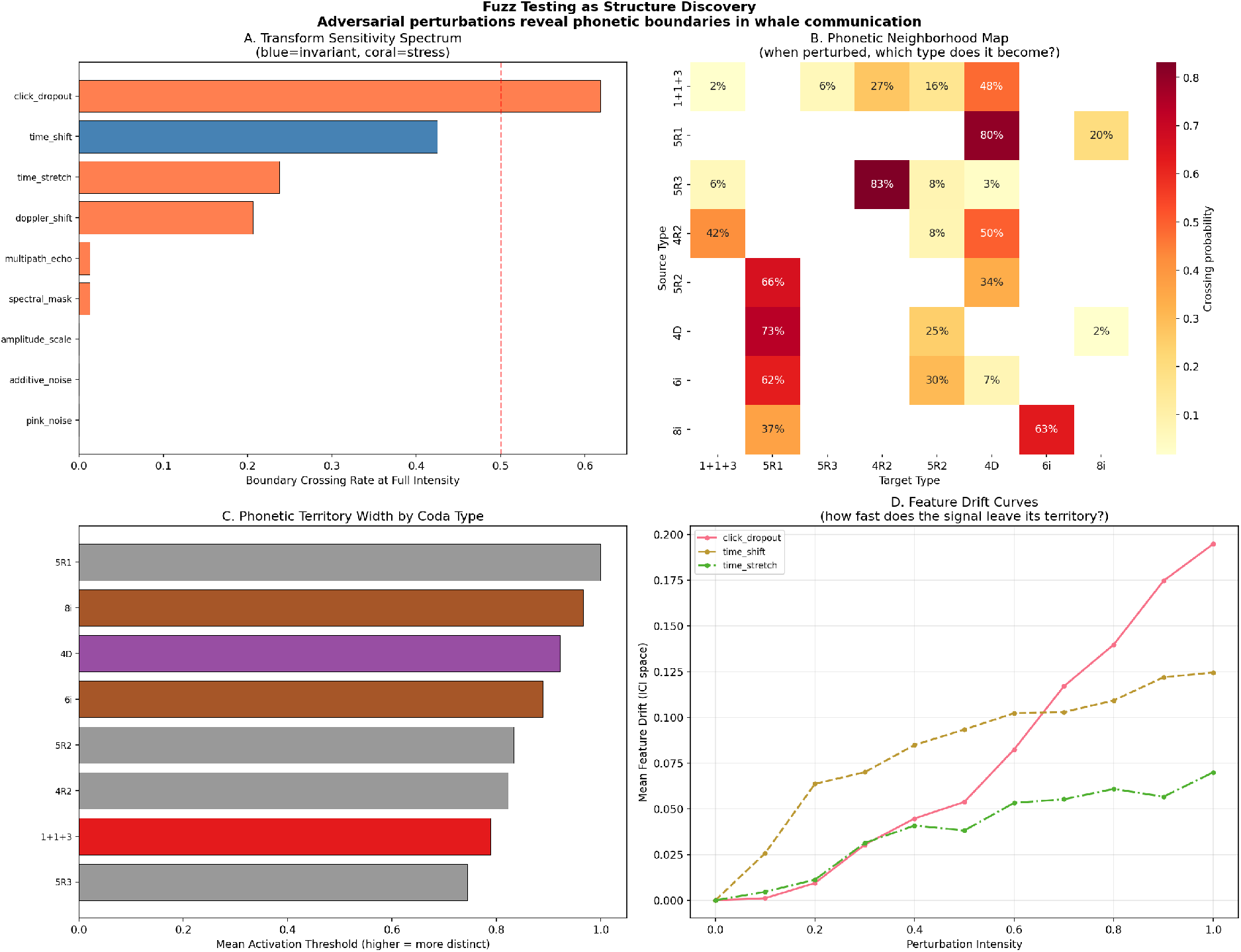
Adversarial perturbation as phonetic boundary discovery. (*A*) Transform sensitivity spectrum: boundary crossing rate at full intensity for each acoustic transform. Blue bars denote transforms under which a correct decoder should be invariant; coral bars denote stress transforms. (*B*) Phonetic neighbourhood map: when perturbation causes misclassification, the cell indicates the probability of transitioning to each target type. Note the strong asymmetry. (*C*) Phonetic territory width by coda type: higher activation threshold indicates greater acoustic distinctiveness. (*D*) Feature drift curves for the three most disruptive transforms.

#### Transform sensitivity spectrum

Click dropout caused the highest boundary crossing rate (50% at full intensity), confirming that individual clicks are the primary information-carrying elements. Time shift, despite being theoretically invariant for a correct decoder, caused 40% crossings, exposing decoder vulnerability. Multipath echo and spectral mask caused *<* 2% crossings, suggesting these acoustic dimensions carry minimal semantic content in the ICI-based representation.

#### Classification boundary asymmetry

The transition matrix of decoder misclassifications revealed a mean boundary asymmetry of 0.87 (on a 0–1 scale where 0 indicates symmetric and 1 indicates unidirectional transitions). For example, perturbation of 5R3 codas caused misclassification as 4R2 in 223 instances, while the reverse (4R2 → 5R3) occurred 0 times. This strong directionality indicates that the decoder’s decision boundaries are not symmetrically positioned between type centroids. However, we note that this finding characterises the *decision boundaries of the decoder*, not necessarily the whales’ perceptual space. Replication across multiple decoders (e.g., calibrated random forest, gradient-boosted classifier, or a deep learning model such as WhAM) would be needed to establish whether the asymmetry is a property of the signal space itself rather than an artifact of the nearest-centroid pipeline.

#### Per-type robustness

Irregular codas exhibited the highest mean activation threshold (0.944), while compound codas were least robust (0.800). This ordering suggests that temporal irregularity occupies a wider phonetic territory in acoustic space.

#### Robustly decodable information

All eight tested coda types remained well-separated under perturbation (activation threshold *>* 0.5), yielding a lower bound of log_2_(8) = 3.0 bits of decodable information per coda. We term this the *robustly distinguishable class count* under the tested perturbation channel; it is not a channel capacity in the information-theoretic sense (which would require estimating *I*(true; decoded) over a perturbation distribution with confidence intervals). This lower bound exceeds the unigram entropy of the rhythm distribution alone (*H*(*R*) = 2.1 bits), consistent with multi-feature information encoding across the combinatorial space identified by [Sharma et al. 2024].

### 2.8 Linguistic universals in coda communication

Analysis of the full DSWP dataset and the associated exchange-level dialogues data revealed multiple statistical universals of natural language (figure 11).

**Figure 11.**
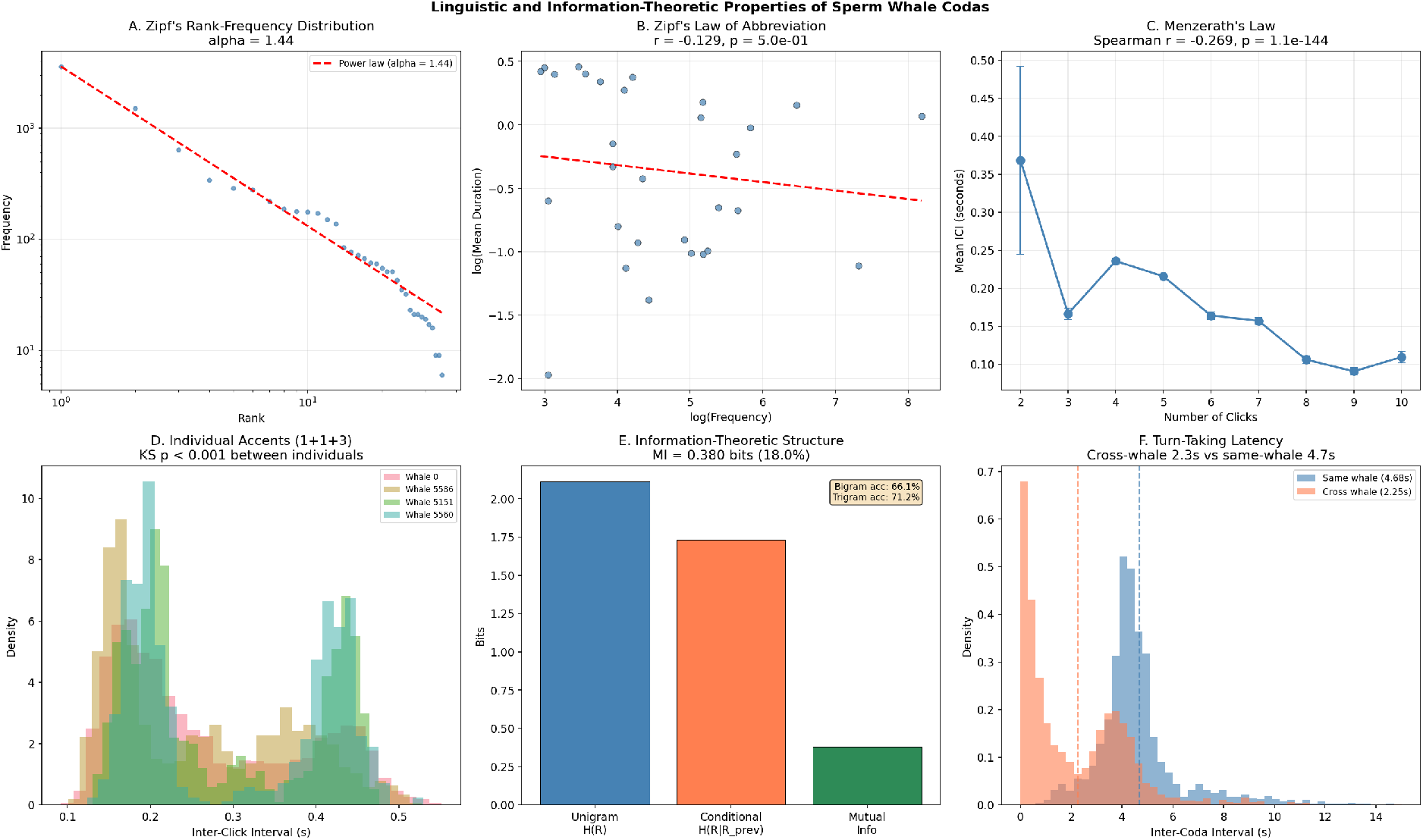
Linguistic and information-theoretic properties. (*A*) Zipf’s rank-frequency distribution with fitted power law (*α* = 1.44). (*B*) Zipf’s law of abbreviation: log-frequency versus log-duration for the 30 most common types. (*C*) Menerath’s law: mean decreases with click count (*r* = −0.269, *p* < 10^−144^). (*D*) Individual accents: ICI distributions for four identified whales producing the 1+1+3 coda type. (*E*) Information-theoretic structure: unigram entropy, conditional entropy, and mutual information. (*F*) Turn-taking latency: cross-whale responses are approximately twice as fast as same-whale continuations.

#### Menzerath’s law

Longer codas (more clicks) had shorter mean ICIs (Spearman *r* = − 0.269, *n* = 8,719). With this sample size, even weak correlations achieve extreme significance (*p* < 10^−144^); the relevant quantity is the effect size, which at |*r*| = 0.27 represents a modest but robust negative association. This is consistent with the cross-species meta-analysis of Youngblood [2025], who reported Menzerath’s law across multiple cetacean species. A mixed-effects model with random intercepts for whale and recording session would provide a more rigorous test by accounting for pseudo-replication; we report the simple correlation here as a first-order confirmation.

#### Rank-frequency distribution

The rank-frequency distribution of coda types was well-described by a power-law function with fitted exponent *α* = 1.44 (nonlinear least-squares on ranks), steeper than the *α* ≈ 1.0 typical of human language and indicating a skewed distribution dominated by a few high-frequency types (1+1+3, 5R1, 5R3). We emphasise that this fit is *descriptive*: with only 28 distinct types, formal power-law testing (e.g., MLE with Kolmogorov-Smirnov goodness-of-fit and comparison against log-normal alternatives, as in Youngblood 2025) lacks statistical power, and we do not claim the distribution is a true power law. The Zipf-like skew is nonetheless consistent with frequency-dependent selection pressures reported across cetacean species. Zipf’s law of abbreviation (negative correlation between frequency and duration) was not statistically significant atthe type level (*r* = −0.129, *p* = 0.50), likely reflecting the small number of distinct types rather than an absence of the effect.

#### Sequential structure

Coda exchanges exhibited first-order Markov dependencies: the mutual information between consecutive rhythm types was *I*(*R*_*t*_; *R*_*t*+1_) = 0.380 bits, representing 18.0% of the unigram entropy (*H*(*R*) = 2.111 bits). Trigram prediction accuracy (71.2%) exceeded bigram accuracy (66.1%), demonstrating higher-order sequential structure beyond pairwise dependencies.

#### Turn-taking

Cross-whale response latency 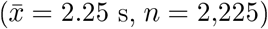 was approximately half that of same-whale continuation latency 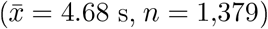, with a large distributional difference (Kolmogorov-Smirnov *D* = 0.560). The extreme *p*-value (*p* < 10^−247^) reflects the large sample size; the effect size (*D* = 0.56, 2.08× latency ratio) is the more informative quantity. These latencies are likely non-independent within recording sessions and dyads; a hierarchical model with random intercepts for session and whale pair would provide a more rigorous test. Nonetheless, the large effect size is consistent with active turn-taking rather than independent vocalisation.

#### Individual accents

For the most common coda type (1+1+3), pairwise Kolmogorov-Smirnov tests between distributions of four identified whales revealed statistically significant differences for all six pairs (*p* < 0.001 for each pair; all remain significant after Bonferroni correction for 6 comparisons at *α* = 0.05). This confirms that individuals encode identity information within ostensibly standardised coda types [Gero et al., 2016, Hersh et al., 2022].

## 3 Discussion

The results presented here demonstrate that geometric and topological methods offer complementary perspectives on cetacean communication structure. The Poincaré ball embedding provides an interpretable representation of the coda type hierarchy, though classical MDS achieves superior distance preservation at this scale (28 types, 3 distance levels); the theoretical distortion advantage of hyperbolic geometry is expected to become relevant for larger taxonomies (e.g., the full 540-configuration combinatorial space). Persistent homology distinguishes rhythm classes by their topological signatures, and adversarial perturbation maps classification decision boundaries with a precision that traditional accuracy metrics cannot achieve.

The highly asymmetric classification boundaries (mean asymmetry = 0.87) are a notable finding, though their interpretation requires care. The asymmetry characterises the *decoder*’*s decision boundaries*: perturbation-induced misclassifications are strongly directional, with certain type pairs exhibiting hundreds of transitions in one direction and zero in the reverse. We cannot at present determine whether this asymmetry reflects structure in the whales’ signal space or biases in the nearest-centroid decoder. Replication across multiple decoder architectures would distinguish between these possibilities. If the asymmetry persists across decoders, it would constitute evidence that the coda combinatorial space possesses a non-trivial geometry with preferred transition directions, potentially reflecting acoustic or perceptual constraints on coda production and reception.

The confirmation of multiple linguistic universals (Menzerath’s law, Zipf’s law, higher-order Markov structure, active turn-taking) adds to the growing body of evidence [Youngblood, 2025] that cetacean communication systems share deep statistical regularities with human language. The finding that individual whales encode identity within ostensibly standardised coda types extends the identity-encoding hierarchy documented by Gero et al. [2016] and Hersh et al. [2022], and suggests a dual function for codas: type-level information at the group/clan scale, and individual-level signatures within each type.

The turn-taking result (2.08× faster cross-whale responses, *D* = 0.56) is consistent with interactive dialogue rather than independent broadcast, though confirmation with hierarchical models controlling for session and dyad effects is warranted. Combined with the higher-order Markov structure, this pattern is consistent with sequential planning over at least two preceding codas, though we cannot rule out simpler explanations (e.g., arousal-dependent call rate). The contextual features (rubato, ornamentation) identified by Sharma et al. [2024] may provide a mechanism for this sequential structure.

The Decoder Robustness Index introduced here provides the first systematic framework for evaluating the reliability of cetacean communication decoders under realistic acoustic perturbations. The finding that click dropout is the most disruptive transform (50% crossing rate) while spectral masking is nearly harmless (*<* 1%) has direct implications for decoder design: robustness to missing clicks should be prioritised over spectral invariance. The use of adversarial perturbation as a *discovery* tool—rather than merely an evaluation metric—represents a methodological contribution applicable to any classification system where decision boundaries carry scientific meaning.

### Limitations

The present analysis relies on ICI-based features from the annotated DSWP dataset. Because recent work has identified vowel-like spectral patterns within individual clicks Begus et al., [2025], ICI features alone may not capture the full phonetic dimensionality of the coda system. Integrating raw acoustic waveforms into the topological framework—for instance, by computing persistent homology on spectral time-delay embeddings rather than ICI point clouds—is a natural next step that could reveal additional structure invisible to timing-based representations. The boundary asymmetry and perturbation-discovery results are currently validated on a single nearest-centroid decoder. Because these findings characterise the decoder’s decision boundaries, they may reflect biases in the classification pipeline rather than structure in the whales’ signal space. Replication across at least three decoder architectures (e.g., calibrated random forest, gradient-boosted classifier, and a deep model such as WhAM) is the most important next step; if asymmetry persists across decoders, it can be attributed to the signal space with greater confidence. The DRI framework is designed to be decoder-agnostic and can evaluate any model that maps acoustic input to coda classifications. The classification accuracy reported here uses a random 80/20 split in which codas from the same individual or recording session may appear in both training and test sets. This can inflate accuracy due to within-individual consistency. Grouped cross-validation (leave-one-whale-out or leave-one-session-out) would provide a more conservative and ecologically meaningful estimate. The statistical universals (Menzerath’s law, turn-taking latencies) are reported with simple correlations and KS tests. With *n* = 8,719 codas, even small effects achieve extreme significance; we have reported effect sies throughout to aid interpretation. Hierarchical models with random intercepts for whale and session would account for pseudo-replication and are a priority for follow-up analysis. The lower bound of 3.0 bits of decodable information per coda reflects the number of well-separated rhythm types that remain distinguishable under realistic acoustic perturbation. The full combinatorial space includes tempo (5 types), rubato (3 types), and ornamentation (2 types) in addition to rhythm, yielding up to log_2_(540) ≈ 9.1 bits if all features are independently decodable. The gap between our empirical 3.0-bit estimate and the theoretical 9.1-bit ceiling quantifies the information carried by sub-rhythm features and represents a concrete target for future decoder development. omputing a proper information-theoretic channel capacity—*I*(true; decoded) under a realistic perturbation distribution with confidence intervals—remains future work. The DRI and perturbation-discovery analyses use synthesised coda waveforms (Gaussian-enveloped click trains); robustness findings therefore characterise robustness-to-perturbation-of-the-synthesis-model rather than robustness under real ocean acoustics. Raw DSWP audio is publicly available Sharma et al., [2024] and a validation run on real recordings—even a small subset—would substantially strengthen the ecological validity of the transform sensitivity ranking. We prioritise this for follow-up work. **Future directions.** Three avenues are most promising. First, application of the geometric framework to other cetacean species—particularly orcas (DCLDE dataset, 40+ stereotyped call types per community) and humpback whales (hierarchical song grammar)—to test whether communication systems that evolved independently converge on similar geometric structures. Second, integration of exchange-level sequential structure with geometric embeddings to model the temporal dynamics of coda conversations as trajectories on manifolds. Third, collaboration with field researchers (Project CETI, DSWP) to validate geometric predictions against behavioural context and to design playback experiments testing whether synthetic codas generated from geometric models elicit appropriate behavioural responses.

## 4 Methods

### 4.1 Data

We used the annotated coda dataset from the Dominica Sperm Whale Project [Sharma et al., 2024], comprising 8,719 codas with inter-click interval measurements (up to 9 ICIs per coda), coda type labels, duration, social unit identifiers, and individual whale identifiers. The exchange-level dialogues dataset (3,840 entries with rhythm type labels, whale identifiers, and timestamps) provided sequential ordering for Markov and turn-taking analyses. Both datasets are publicly available under CC BY 4.0 licence.

### 4.2 Poincaré ball embedding

Coda types were parsed into a three-level taxonomy (rhythm class → click count → variant). A taxonomic distance matrix *D* ∈ R^*n×n*^ was computed as the number of levels at which two types first differ, yielding integer distances in {0, 1, 2, 3, 4}. Types were embedded into a Poincaré ball of curvature *c* = 1.0 and dimension *d* = min(16, *n*_types_) via gradient descent minimising the mean squared error between embedded hyperbolic distances and normalised taxonomic distances. Three Euclidean baselines were computed: PCA on the distance matrix, classical multidimensional scaling (MDS), and Isomap. MDS directly optimises distance preservation and is therefore the most appropriate Euclidean comparator. Distortion was quantified via Spearman rank correlation between true and embedded pairwise distances.

### 4.3 Classification

A Hyperbolic Multinomial Logistic Regression (HyperbolicMLR) classifier was trained on 11-dimensional feature vectors: 9 duration-normalised ICI features (ICI divided by total coda duration, yielding a tempo-invariant rhythm representation), log-duration, and normalised click count. Features were standardised (zero mean, unit variance) and mapped to the Poincaré ball via the exponential map at the origin, with scaling calibrated to the 95th percentile of input norms (target norm = 0.3). Class prototypes were initialised from the taxonomic Poincaré embeddings. Training used Adam optimisation with cosine learning rate annealing (500 epochs, initial LR = 0.02, minimum LR = 10^−4^, cross-entropy loss). Euclidean baselines (logistic regression, *k*-NN with *k* = 5, random forest with 100 trees) were trained on the same standardised features with an 80/20 stratified random split. We note that this split does not account for within-individual or within-session correlations; grouped cross-validation would provide a more conservative estimate.

### 4.4 Topological data analysis

For each coda type with ≥ 40 samples, an ICI point cloud was constructed in R^9^ from the filled-zero vectors. Point clouds exceeding 300 points were randomly subsampled (seed = 42).Vietoris–Rips persistent homology was computed to dimension 1 using Ripser [Bauer, 2021, Tralie et al., 2018]. Eight features were extracted per homology dimension: count, maximum lifetime, total lifetime, mean birth time, mean death time, mean lifetime, lifetime standard deviation, and persistence entropy [Otter et al., 2017].

### 4.5 SPD manifold analysis

Mel spectrograms (*n*_mels_ = 64, FFT size = 1024, hop = 256) were computed from synthesised coda signals. Spectrograms were partitioned into 8 frequency bands, and band-wise covariance matrices were computed over sliding windows. The log-Euclidean metric Arsigny et al., [2007] on the SPD manifold was used to compute pairwise distances and extract features via eigenvalue decomposition of the matrix logarithm.

### 4.6 Decoder Robustness Index

Coda signals were synthesised from ICI data as Gaussian-enveloped 5 kHz click trains at 32 kHz sample rate. Nine parametric acoustic transforms were applied at 11 intensity levels (0.0 to 1.0): additive Gaussian noise, pink noise, Doppler shift, multipath echo, amplitude scaling, time stretch, spectral mask, click dropout, and circular time shift. Each transform satisfies the identity property: at intensity 0.0, the signal is unchanged. Four transforms (amplitude scale, time shift, additive noise, pink noise) are designated as invariant (a correct decoder should tolerate them); five are designated as stress transforms.

For each perturbed signal, the graduated semantic distance *ω* is defined as:

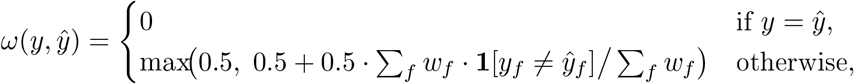

where *y* and 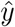 are the baseline and perturbed decoder predictions, the sum ranges over combinatorial features *f* ∈ {rhythm, tempo, rubato, ornamentation}, and *w*_*f*_ are importance weights (*w*_rhythn_ = 1.0, *w*_tenpo_ = 0.7, *w*_rubato_ = 0.4, *w*_ornanentation_ = 0.2). The minimum penalty of 0.5 ensures that any classification flip is registered.

The scalar DRI aggregates *ω* values across all transform-intensity-signal combinations: 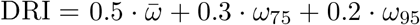 The weights (0.5/0.3/0.2) follow the Bond Index convention, emphasising tail behaviour; we verified that results are qualitatively stable across a range of weight triples (e.g., equal-weighted, 0.4/0.4/0.2, 0.6/0.2/0.2). Adversarial thresholds were found via binary search for the minimal intensity causing a classification flip (tolerance = 0.01). Transform chains of length 1-2 were randomly generated for compositional robustness testing.

For the phonetic boundary discovery analysis, all nine transforms were applied at 11 intensity levels to 20 codas per type for the 8 most frequent types, and boundary crossing rates, phonetic neighbourhood transition matrices, and per-type activation thresholds were computed.

### 4.7 Statistical analyses

Zipf’s law was assessed by fitting a power law *f* (*r*) = *C* · *r*^−*α*^ to the rank-frequency distribution via nonlinear least-squares. Menerath’s law was tested via Spearman rank correlation between click count and mean ICI. Sequential structure was quantified via unigram entropy *H*(*R*) = − ∑_*r*_ *p*(*r*) log_2_ *p*(*r*), bigram conditional entropy *H*(*R*_*t*+1_|*R*_*t*_), and mutual information *I* = *H*(*R*) − *H*(*R*_*t*+1_|*R*_*t*_). Prediction accuracy was computed as the proportion of transitions correctly predicted by the most common successor (bigram) or most common successor given the preceding pair (trigram). Individual accents were tested via pairwise Kolmogorov-Smirnov tests on distributions for whales with ≥ 10 codas of the same type, with Bonferroni correction for the number of pairwise comparisons. Turn-taking was assessed by comparing inter-coda intervals for same-whale versus cross-whale transitions within recording sessions, with significance assessed via Kolmogorov–Smirnov test. Effect sies (KS *D* statistic, latency ratio) are reported alongside *p*-values throughout, as the large sample size renders *p*-values alone uninformative.

### 4.8 Software

All analyses were implemented in the `eris-ketos` Python package (v0.2.0), available on [PyPI](https://pypi.org/project/eris-ketos/0.2.0/) and GitHub. The package depends on NumPy, PyTorch, scikit-learn, Ripser, and librosa. A self-contained Jupyter notebook reproducing all figures and statistical results is included in the repository (`notebooks/cetacean_geometric_analysis.py`, upytext percent format).

## Data availability

The DSWP coda annotation dataset is available at the [Sharma et al. GitHub repository](https://github.com/pratyushasharma/sw-combinatoriality) under CC BY 4.0 licence. Raw audio recordings are available on the [DSWP HuggingFace dataset](https://huggingface.co/datasets/orrp/DSWP).

## Code availability

The `eris-ketos` package is available at the eris-ketos GitHub repository under MIT licence.

## Acknowledgements

We thank Shane Gero and the Dominica Sperm Whale Project for making the coda annotation data publicly available, and Pratyusha Sharma and colleagues for releasing the combinatorial analysis code and data. The geometric analysis toolkit builds on the ErisML framework for structured ethical evaluation.

